# SHIP2 controls matrix mineralization by regulation of the RhoA/ROCK pathway and remodeling of the actin cytoskeleton

**DOI:** 10.1101/2022.10.30.514432

**Authors:** Anais Fradet, Jamie Fitzgerald

## Abstract

Mutations in *INPPL1*, the gene coding for SH2 Domain-Containing Inositol 5’-Phosphatase 2 (SHIP2), cause Opsismodysplasia, a severe chondrodysplasia characterized by delayed bone maturation. The mechanism by which the loss of an inositol phosphatase causes a major skeletal developmental defect is unclear. To investigate the role of SHIP2 in mineralization, the *INPPL1* gene was deleted *in vitro* in chondrocyte and osteoblast differentiation models and the effect of the loss of SHIP2 on cell differentiation, subsequent mineralization, and on actin cytoskeleton formation was investigated. The loss of SHIP2 does not impact differentiation but, consistent with the disease phenotype, induces a significant reduction in extracellular matrix mineralization in both cell types. Absence of SHIP2 also altered the actin cytoskeleton to increase cell adhesion and focal adhesion formation. Furthermore, inhibition of actin polymerization in SHIP2-deficient cells rescued the mineralization phenotype. RhoA/ROCK, Cdc42 and Rac1 are the three main RhoGTPases responsible for actin cytoskeleton regulation in bone cells. Specific inhibitors of these RhoGTPases were used to determine the pathways involved in SHIP2-mediated mineralization. Since only the ROCK pathway inhibitor rescued the mineralization phenotype, it is concluded that SHIP2 regulates actin cytoskeleton remodeling and consequently extracellular matrix mineralization by inhibiting the RhoA/ROCK pathway.

## Introduction

The protein SHIP2, encoded by the *INPPL1* gene, is an inositol polyphosphate 5-phosphatase. Its main function is to dephosphorylate the lipid second messenger phosphatidylinositol 3,4,5-triphosphate (PI(3,4,5)P3) to phosphatidylinositol 3,4-diphosphate (PI(3,4)P2). In addition to its catalytic domain, SHIP2 contains several motifs known to be involved in protein-protein interactions and has been shown to function as a docking protein for a variety of intracellular molecules. These include an N-terminal SRC homology 2 (SH2) domain, potential phosphotyrosine binding sites containing the consensus NPxY, a sterile alpha motif (SAM) domain and C-terminal proline-rich domains with potential for SH3 binding. SHIP2 is prominent in a range of human diseases with most research focused on diabetes, cancer and more recently Alzheimer’s disease (for reviews see (1–3)). In 2013 it was reported that compound heterozygous mutations in *INPPL1* caused Opsismodysplasia, a rare skeletal chondrodysplasia primarily characterized by growth plate defects and delayed bone maturation (4–6). While the precise disease mechanisms are unknown, the majority of mutations described so far are predicted to abolish SHIP2 catalytic activity (7,8). *In vivo* studies suggest a role for SHIP2 in a variety of processes including fibroblast growth factor signaling, mitogen-activated protein kinase (MAPK) signaling and phosphoinositide–kinase (PI3K) signaling (9–12) and as an important regulator of insulin signaling via the Akt pathway (13).

SHIP2 is also involved in cell adhesion and cell polarity regulation. SHIP2 is a member of the adhesome, a theoretical network that has been assembled through the analysis of proteins that localize to integrin adhesion complexes (14). Remodeling of the actin cytoskeleton through actin dynamics is important in several biological processes including cell adhesion, cell migration, and cell morphology. These processes are known to regulate chondrocyte differentiation and more generally endochondral ossification (15,16). Importantly, several human chondrodysplasias have been linked to mutations affecting the actin cytoskeleton (17).

Rho GTPases control cell adhesion and migration processes via regulation of cytoskeletal assembly and maintenance (18). In mammals, the Rho GTPase family contains 20 members with most research focused on Cdc42, Rac1 and RhoA. RhoA, through ROCK, plays a prominent role in endochondral ossification. In osteoblasts, RhoA/ROCK negatively regulates BMP signaling. Moreover, expression of a dominant-negative ROCK mutant in mouse stromal ST2 cells induced osteoblastic differentiation, whereas expression of a constitutively active ROCK mutant attenuated osteoblastic differentiation (19,20). Consistent with these findings, activation of RhoA/ROCK with a toxin almost completely abolishes osteoblast differentiation of calvaria cells (21).

Mature chondrocytes display a characteristic cortical actin organization while the organization of actin stress fibers has been associated with de-differentiation of articular chondrocytes *in vitro* (22,23). RhoA controls the formation of actin stress fibers in numerous cells (24) and in chondrocytes, overexpression of RhoA inhibits both early chondrogenesis and hypertrophic chondrocyte differentiation, while inhibition of RhoA/ROCK promotes chondrocyte maturation (22,25). The effects of RhoA/ROCK signaling on chondrogenesis seems to be mediated at least in part by regulation of Sox9, the central transcriptional regulator of chondrogenesis (22,26,27). In glioma cells, SHIP2 and RhoA have been shown interacting in a GTP manner. In these cells, depletion of SHIP2 leads to a defect in cell polarization and cell migration (28).

In the present study, the relationship between SHIP2, RhoA and mineralization in the two major cell types responsible for skeletal mineralization; chondrocytes and osteoblasts was studied. Evidence is provided that SHIP2 inhibits the RhoA/ROCK pathway in both chondrocytes and osteoblasts *in vitro,* resulting in changes to the actin cytoskeleton and inhibition of mineralization.

## Materials and Methods

### Cell culture

ATDC5 cells (Sigma-Aldrich #99072806 supplied by E-ACC) were cultured in DMEM/F12 (Gibco) supplemented with 1 % GlutaMax (Gibco), 10 % fetal bovine serum (FBS, Gibco) and 1 % Antibiotic-Antimycotic (Gibco). Chondrogenic differentiation was conducted as described (29). Briefly, ATDC5 were seeded at 6×10^3^/ cm^2^ in 24-well plates and cultured in DMEM/F12 supplemented with 1 % GlutaMax, 5 % FBS, 1 % Insulin Transferin Selenium (ITS, Gibco) and 1 % Antibiotic-Antimycotic for 5 d until confluency. To induce mineralization, the media was then supplemented with 10 mM β-glycerophosphate (Sigma-Aldrich) and 50 ug/mL of L-Ascorbic acid 2-phosphate (Sigma-Aldrich) for a total of 28 d. SaOs-2 cells (ATCC® HTB-85™) were cultured in McCoy media (Gibco) supplemented with 10 % FBS and 1 % Antibiotic-Antimycotic. For osteogenic differentiation, the media was supplemented with 50 ug/mL of L-Ascorbic acid 2-phosphate and 7.5 mM of B-glycerophosphate for 14 days.

### Cell staining

After differentiation, 3 wells were used for each staining. For Alizarin Red S staining, cells were fixed with 4 % PFA for 5 min at 4 °C and then stained with 500 μL of 2 % Alizarin red S at pH 4.2 5 min at room temperature. Staining was then extracted with 500 μL of 10 % cetylpyridinium chloride 10 min at room temperature and absorbance was measured at 570 nm. For Alcian Blue staining, cells were fixed with 95 % methanol for 20 min at room temperature and then stained with 500 μL of 1 % Alcian blue 8GX in 0.1 M HCl overnight at room temperature. Staining was then extracted with 1 mL per well of guanidine-HCl 6 M 6 h at room temperature and absorbance was measured at 630 nm. Results are representative of three independent experiments with triplicate wells per experiment.

### Deletion of SHIP2

ATDC5 and SaOS-2 cells were transfected with the mouse or human SHIP2 double nickase plasmid (Santa Cruz, sc-421138-NIC (mouse), sc-401622-NIC (human)) using the UltraCruz transfection reagent and the Plasmid transfection medium (Santa Cruz) following manufacturer’s instructions. Briefly, 2×10^5^ cells were transfected using 1 μg of plasmid and 10 μl of transfection reagent. After transfection GFP positive cells were sorted by flow cytometry and further selected in 1.5 μg/ml puromycin. Selected clones were then screened by Western Blot. Two clones per cell line were kept for further experiments.

### Western Blot

Cell proteins were extracted using a NP-40 lysis buffer (150 mM sodium chloride; 1.0 % NP-40; 50 mM Tris pH 8.0) and protein concentration determined using a BCA assay (Pierce). The same amount of proteins for each lysate was then separated by SDS-PAGE on 4-15 % precast gels (Bio-Rad) under reducing conditions, and transferred to PVDF membranes (Millipore) using a wet transfer system. After transfer, the membrane was blocked using the Odyssey blocking buffer (LiCor) and incubated overnight at 4 °C with the primary antibody (SHIP2 antibody (Abcam #ab70267) 1/2000, MMP13 antibody (Novus #NBP1-45723SS) 1/500, Akt antibody (R&D system #AF1775) 1/2000, pAkt S473 antibody (Cell Signaling Technologies #4051) 1/500, Erk1/2 antibody (Cell Signaling Technologies #9107) 1/2000, pErk1/2 Thr202/Tyr204 (Cell Signaling Technologies #9101) 1/1000) followed by an 1 h incubation with an IRD800-conjugated goat anti-rabbit or anti-mouse antibody (LiCor, dilution 1/20000). Bands were revealed using a LiCor Odyssey scanner. Tubulin or Total protein staining (LiCor) was used as a loading control. Band intensity was quantitated using the LiCor software.

### Real-time QPCR

Total RNA was extracted from cells using TriReagent (Sigma). 1 μg of RNA was then reverse transcribed into cDNA using iScript kit from Bio-Rad. cDNAs were diluted 1/20 prior amplification. Real-time QPCR was carried out using SYBR Green Mastermix (Applied Biosystems) with specific primers whose efficiency had been previously checked. Data analysis was carried out using the comparative ΔCt method in triplicates. HPRT1 and PPIA were used as housekeeping genes as previously described for ATDC5 and Saos-2 cells (30,31). Results are representative of three independent RNA extractions.

### Zymography

25 μg of proteins from serum-free phenol red-free conditioned media were separated by SDS PAGE using a 10 % gel containing 0.15 mg/ml of Collagen I (Sigma-Aldrich) under non reducing conditions. The gel was then incubated in developing buffer (50 mM Tris, 10 mM CaCl_2_, 50 mM NaCl, 0.05 % Brij 35 (Sigma-Aldrich), pH 7.6) at 37 °C for 48 h. After incubation, gels were stained with Coomassie Blue R-250 and then destained. Collagenolytic activities were detected as clear bands against a Coomassie Blue–stained gel background. Results are representative of two independent experiments.

### Immunofluorescence

Cells were grown on glass coverslip for 48 h and then fixed with 4 % paraformaldehyde in PBS for 10 min and permeabilized with 0.1 % Triton X-100 in PBS. Immunodetection was carried out using a mouse polyclonal antibody against Vinculin at 1/50 (Santa Cruz) at 4 °C overnight and the secondary antibody (Cy3-conjugated goat anti-mouse; Abcam). The distribution of F-actin was visualized using phalloidin coupled to FITC. For PI(3,4,5)P3 staining, the mouse antibody from Echelon Biosciences (Z-P345B) was used using the conditions recommended by the manufacturer. Briefly, 0.1 % Saponin was used for cell permeabilization and the primary antibody diluted at 1/100 was incubated for 1 h. Focal adhesion number and surface were measured on four different fields per clone using Image J software as described by Horzum et al. (32).

### Adhesion assay

Plates were coated with fibronectin (1 μg/cm_2_). Cells were seeded at 8×10_4_ cell/ cm_2_ and incubated 30 min at 37 °C. Cells were then fixed with 96 % ethanol and stained with 0.1 % crystal violet for 30 min. Finally, cells were permeabilized with 0.2 % Triton overnight and absorbance was read at 595 nm. Results are representative of three independent tests with n=10 wells for each clone in each assay.

### Statistical analysis

Data are expressed as mean ± SD. Statistically significant differences between WT cells and SHIP2-negative cells were assessed using Student’s t test (two-tailed) when variances were equal and Welch’s t test (two-tailed) when variances were not equal.

## Results

The ATDC5 murine cell line and the SaOs-2 human osteosarcoma cell line, commonly used as models for chondrocyte and osteoblast differentiation, respectively (33–35), were used to study the role of SHIP2 in endochondral ossification *in vitro.* To investigate mineralization in these models, two approaches were employed; genetic deletion of the gene encoding SHIP2, *Inppl1,* using the CRISPR/Cas9 system and pharmacological inhibition of SHIP2 catalytic activity using AS1949490. ATDC5 and Saos-2 cells were deleted for *INPPL1* and the absence of SHIP2 protein confirmed by immunoblotting cell extracts from two ATDC and Saos-2 independent clones (Figs. 1a and b). In mineralization assays, alizarin red staining was reduced in the matrix of ATDC5 and SaOs-2 clones deficient for SHIP2 (Fig. 1c and d). Similarly, when wild-type ATDC5 and Saos-2 cells were differentiated in the presence of a specific SHIP-2 inhibitor, mineralization was markedly reduced (Figs. 1e and f). These *in vitro* data are consistent with the human disease phenotype where reduced or absent SHIP2 leads to a mineralization defect.

**Figure1:**
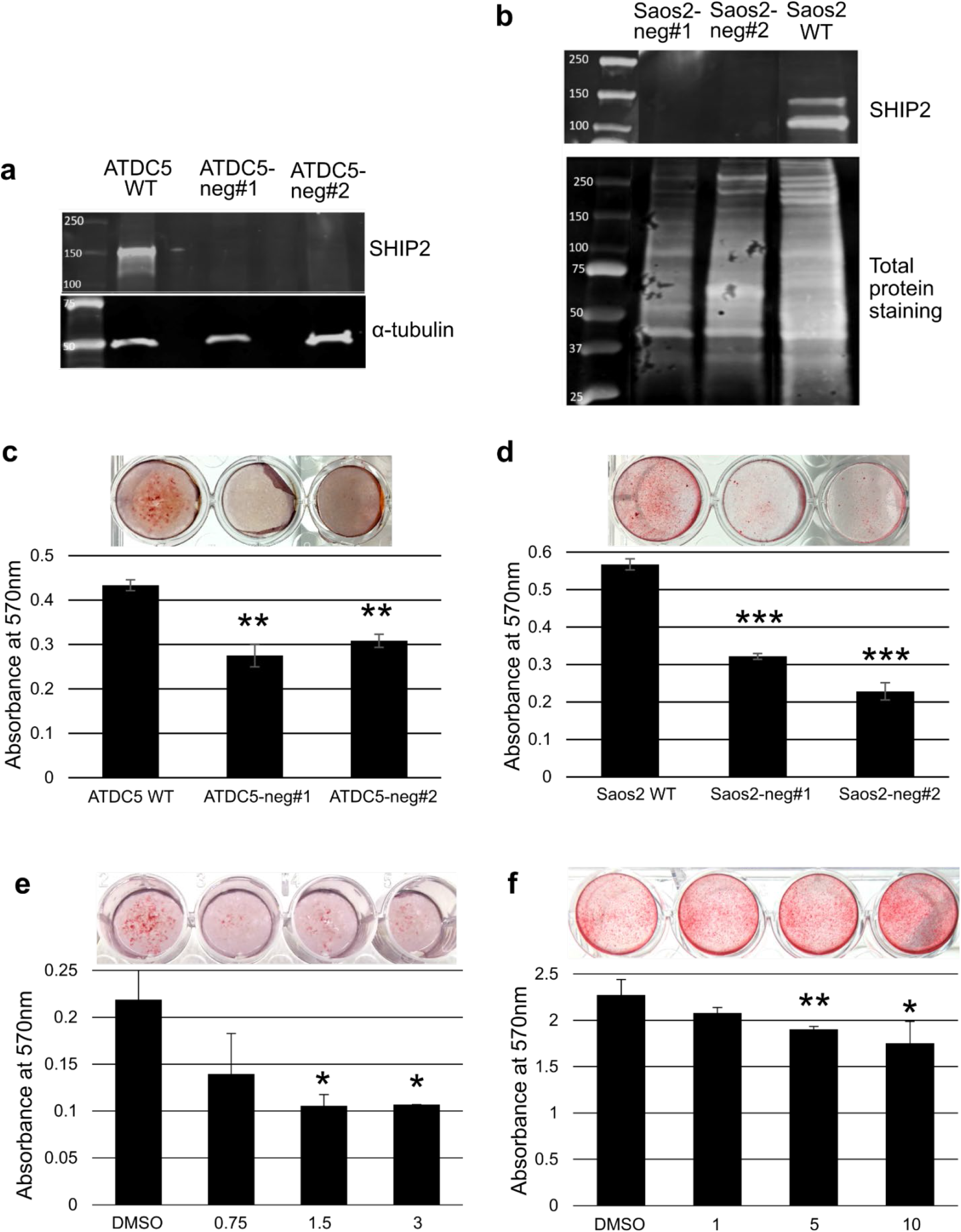
Mineralization defect in chondrocytes and osteoblasts with compromised SHIP2 expression. Immunoblots confirming the absence of SHIP2 protein in independent ATDC5 (a) and Saos-2 clones (b). Wild-type cells plus two clones of ATDC5 (c) and SaOs-2 (d) genetically-depleted for SHIP2 differentiated and then stained with Alizarin Red S. ATDC5 (e) or SaOs-2 (f) cells differentiated in presence of DMSO (vehicle) or SHIP2 inhibitor AS1949490 (0-10 μM) and stained for mineralization with Alizarin Red S. Histograms represent the quantification of the cell staining after extraction. Data is representative of 3 individual experiments (3 wells per experiment). \**p*<0.05, \*\**p*<0.01, \*\*\**p*<0.001.

Associated with reduced mineralization is the finding that differentiated SHIP2-negative ATDC5 cells produce more proteoglycans compared to wild-type cells (Fig. 2a). To explore the cellular phenotype caused by loss of SHIP2 further, the expression of chondrocyte and osteoblast differentiation markers was measured by RT-QPCR in cells following 28 days of differentiation. Compared to wild-type ATDC5 cells, SHIP2-negative cells show a higher expression of *Runx2, Col10a1* (collagen X), *Mmp13, Ocn* (osteocalcin) and *Col1a1* (alpha 1 subunit of collagen I) mRNA (Fig. 2b) demonstrating that these cells approximate the hypertrophic stage of differentiation. The expression of ECM components; *Col2a1* (type II collagen), *Col9a1* (alpha 1 subunit of collagen IX), *Col11a1* (alpha 1 subunit of collagen XI), *Acan* (aggrecan), *Bgn* (biglycan) and *Hspg2* (perlecan), were also upregulated in cells depleted for SHIP2 compared to wild-type ATDC5 chondrocytes (Fig. 2c). In SHIP2-negative SaOs-2 cells, differentiation is demonstrated by the expression of *Runx2, OPN* (osteopontin), *OCN* (osteocalcin), *OSX* (osterix) and *ALP* (alkaline phosphatase) (Fig. 2d). While the expression of these markers increased in SHIP2-negative cells compared to wild-type cells, the expression of *COL1A1* is slightly decreased. Altogether, those results demonstrate that SHIP2 plays a role in the production and mineralization of the matrix but not in the differentiation of either cell type. Confirming the increased Mmp13 mRNA expression seen in Fig. 2b, is the finding by immunoblot and zymography analyses that the level of MMP13 protein (Fig. 3a) and activity (Fig. 3b) is increased in SHIP2-negative ATDC5 cells, compared to WT cells.

**Figure 2:**
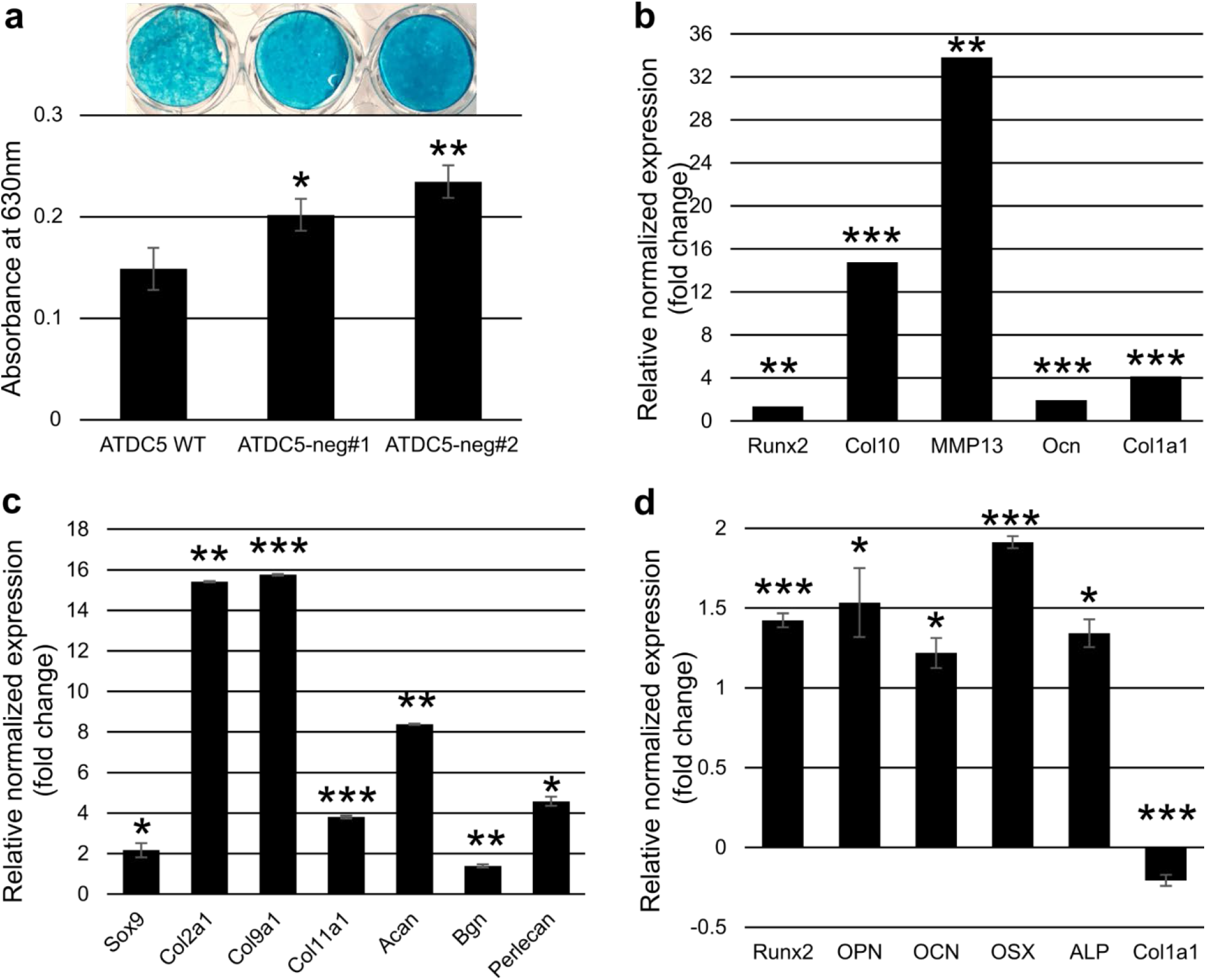
Extracellular matrix production and differentiation master gene expression in SHIP2-negative mineralizing cells. (a) Differentiated ATDC5 were stained for proteoglycans using Alcian Blue. Histograms represent the quantification of the cell staining after extraction. (b) Relative expression of hypertrophy and terminal differentiation markers (c) and matrix related genes in ATDC5-neg#1 cells. (d) Relative expression of osteoblast differentiation markers in SaOs-2 neg#1 cells. Histograms represent the difference in expression compared to WT cells (b, c, d). Data is representative of 3 individual experiments (3 wells per experiment). *p<0.05, **p<0.01, ***p<0.001. WT vs SHIP2-neg.

**Figure 3:**
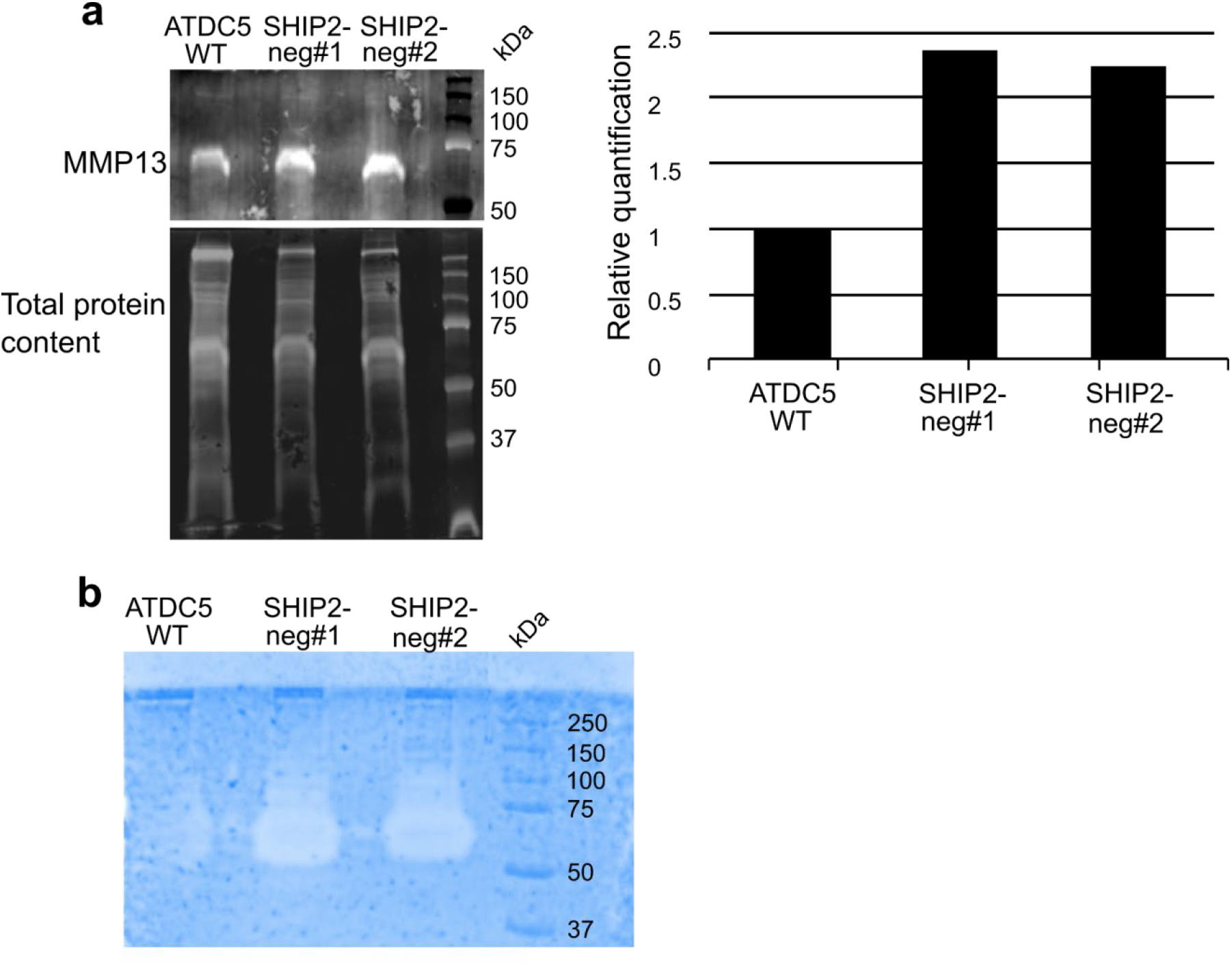
Upregulation of MMP13 in chondrocytes in the absence of SHIP2. (a) Immuno-detection and quantification of MMP13 protein and (b) detection of MMP13 activity by collagen zymography in conditioned media of ATDC5 WT and SHIP2-negative ATDC5 cells. Data is representative of two individual experiments.

One of the major enzymatic functions of SHIP2 is to dephosphorylate PI(3,4,5)P3 at the 5P position to produce PI(3,4)P2. Consistent with this, an accumulation of PI(3,4,5)P3 was observed when ATDC5 cells were treated with the SHIP2 inhibitor AS1949490 or deleted for SHIP2 (Fig. 4a). Surprisingly a reduction in PI(3,4)P2 levels was not observed after inhibition of SHIP2 in ATDC5 by immunofluorescent staining (not shown) possibly reflecting a compensatory production of PI(3,4)P2 through PI3k or PI4k (36). Since PI(3,4,5)P3 is a major second messenger in the Akt pathway, the phosphorylation state of Akt in SHIP2-treated ATDC5 cells was assessed. No increase of Akt phosphorylation was observed in these cells (Fig. 4b).

**Figure 4:**
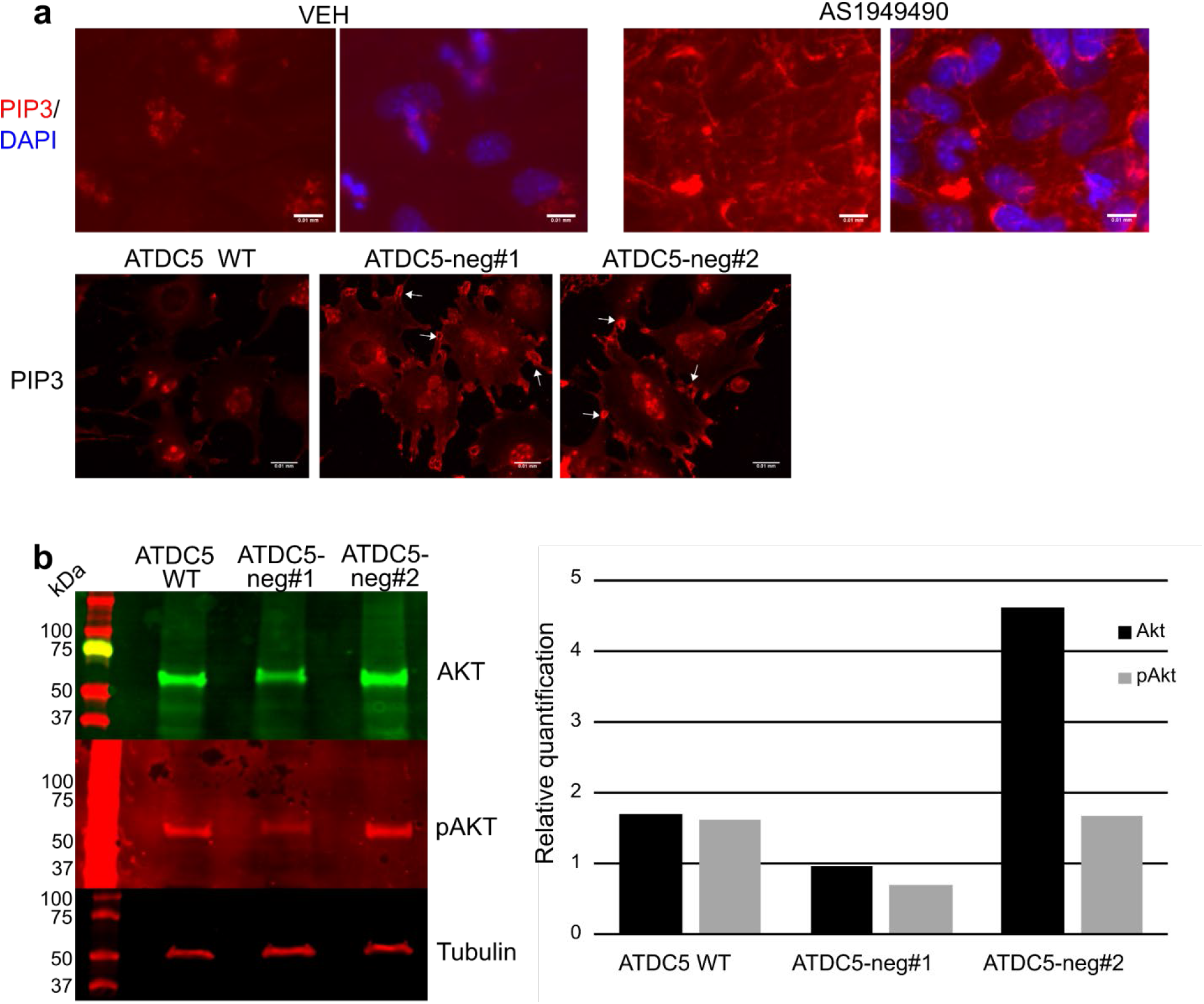
Absence of SHIP2 in chondrocytes leads to the accumulation of PI(3,4,5)P3 without affecting AKT phosphorylation. (a) ATDC5 cells differentiated in presence of DMSO (VEH) or 1.5 μM of AS1949490 (top panels) and SHIP2-negative ATDC5 cells (bottom panels) were immunostained for PI(3,4,5)P3. Arrows indicate focal adhesion-like structures. (b) Immunodetection of Akt and pAKT (S473) protein in SHIP2-negative ATDC5 cells. Scale bars represent 0.01mm.

Interestingly, in SHIP2-negative cells, PI(3,4,5)P3 appeared to localize preferentially in focal adhesion-like structures at the periphery of the cells (arrows in 4a). To determine whether PI(3,4,5)P3 accumulates in focal adhesions, cells were immunostained for the focal adhesion component, vinculin (Fig. 5a). It was not possible to co-stain PI(3,4,5)P3 and vinculin. Nevertheless, vinculin appeared to stain similar structures to PI(3,4,5)P3 in SHIP2-negative ATDC5 cells suggesting a role for SHIP2 in focal adhesion. Moreover, in the absence of SHIP2 the number and the surface area of focal adhesions is increased in ATDC5 and SaOs-2 cells (table, Fig. 5a) and these cells also appeared to be more adherent when plated on fibronectin (Fig. 5b) suggesting that SHIP2 plays a role in regulation of adhesion processes in chondrocytes and osteoblasts.

**Figure 5:**
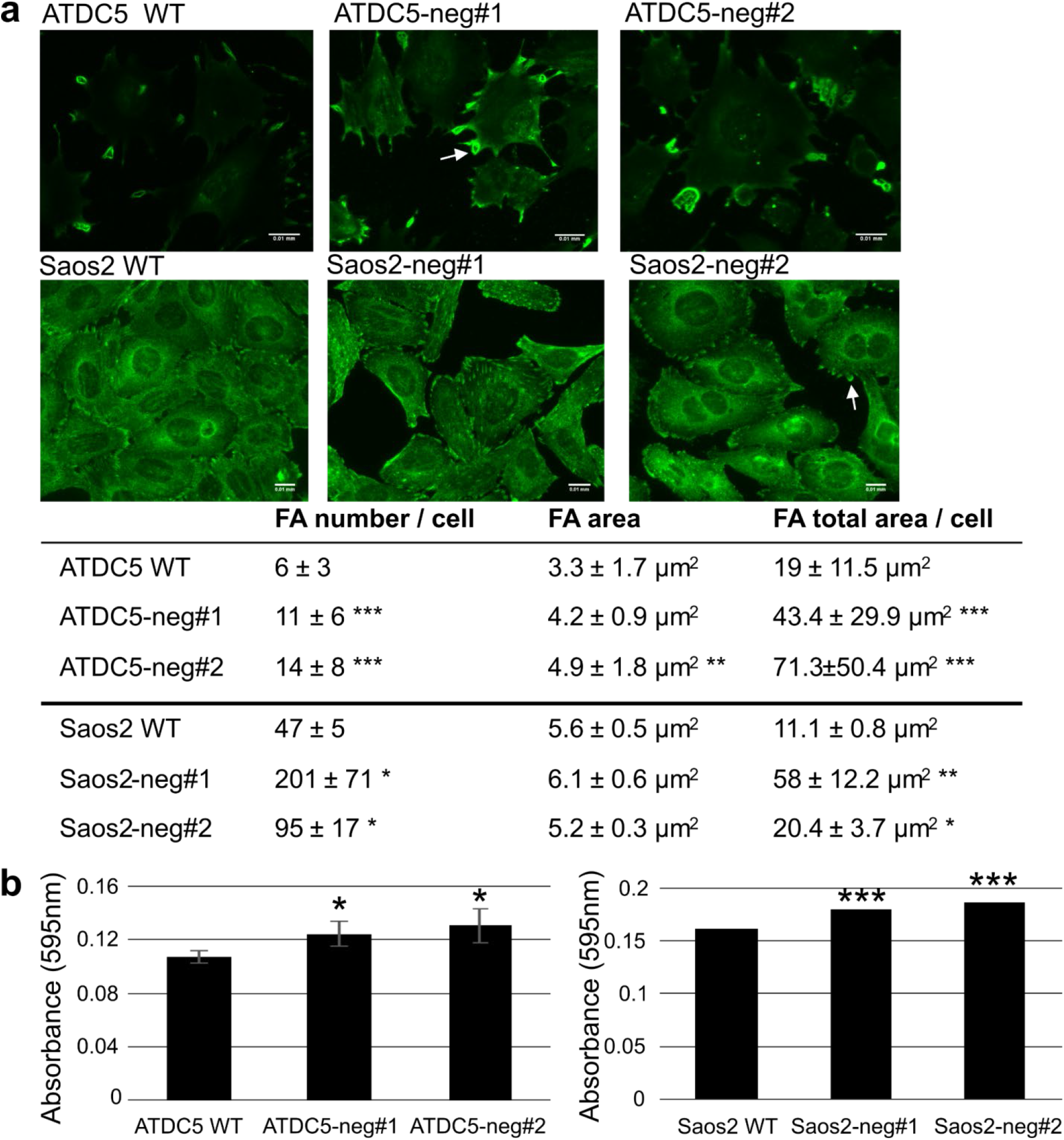
Cells deleted for SHIP2 show numerous larger focal adhesions and have increased adhesion of fibronectin. (a) Wild-type and SHIP2-negative ATDC5 (top panels) and SaOs-2 (bottom panels) cells immunostained for vinculin. Arrows indicate focal adhesion-like structures. Table shows the number of focal adhesions per cell and their surface. (b) Cell adhesion assay showing increased ATDC5 and SaOs-2 attachment 30 min after seeding cells on fibronectin \**p*<0.05, \*\**p*<0.01, \*\*\**p*<0.001 for wild-type vs SHIP2-deleted cells. Scale bars represent 0.01 mm.

Since cell adhesion processes are tightly linked to the modulation of cytoskeleton, cytochalasin D, an inhibitor of actin polymerization, was incubated with ATDC5 chondrocytes and SaOs-2 osteoblasts deleted for SHIP2. Cytochalasin D treatment rescued mineralization in both ATDC5 and SaOs-2 cells depleted for SHIP2 (Fig. 6a). In many cell types including bone cells, actin polymerization is primarily regulated by three small RhoGTPases; RhoA through ROCK, CDC42 and Rac1. To investigate whether these RhoGTPases are involved in the SHIP2-mineralization pathway, differentiated cells were treated with specific inhibitors of these RhoGTPases. Inhibition of Rac1 and Cdc42 did not have any effect on the cells (not shown). However, inhibition of ROCK using Y-27632, an ATP-competitive inhibitor, rescued mineralization in SHIP2-negative cells (Fig. 6b) while partially reducing the synthesis of proteoglycan in ATDC5 cells (Fig. 6c). These data suggest that the ROCK pathway is involved in the inhibition of matrix mineralization induced by the absence of SHIP2.

**Figure 6:**
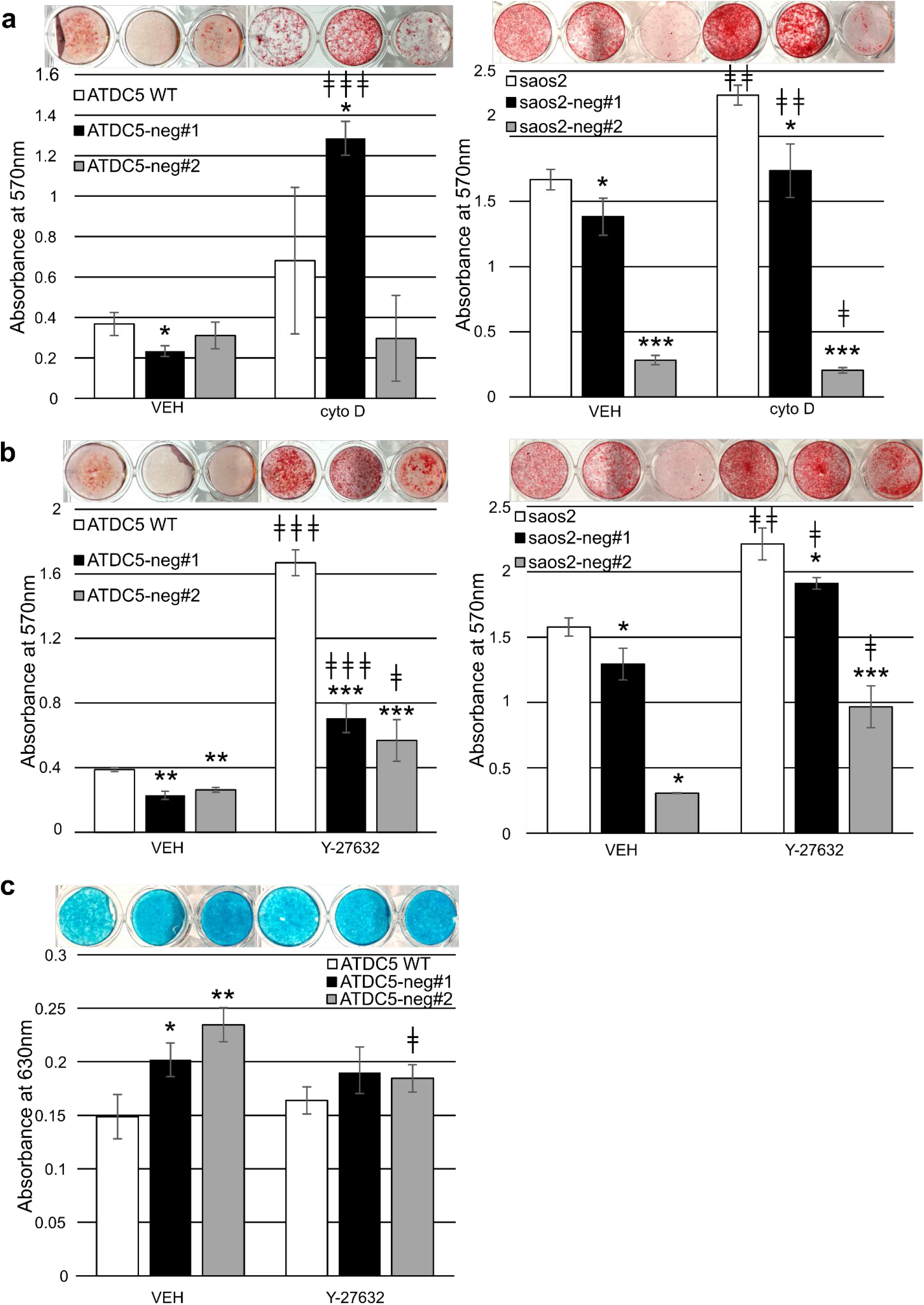
Mineralization phenotype is rescued by actin polymerization inhibitor and ROCK pathway inhibitor. (a) Alizarin Red staining of ATDC5 and SaOs-2 WT cells vs SHIP2-negative cells differentiated in presence of 0.3μM of the actin polymerization inhibitor, cytochalasin D. (b) Alizarin Red staining of ATDC5 and SaOs-2 WT cells vs SHIP2-negative ATDC5 cells differentiated in presence of 15 μM of the ROCK pathway inhibitor, Y-27632. (c) Alcian Blue staining of ATDC5 WT cells vs SHIP2-negative cells differentiated in presence of 15 μM of the ROCK pathway inhibitor, Y-27632. Histograms represent the quantification of the cell staining after extraction. *,‡*p*<0.05, **,‡‡*p*<0.01, ***,‡‡‡*p*<0.001. * WT vs SHIP2-neg. ‡ vehicle vs treatment.

## Discussion

Mutations in *Inppl1,* the gene encoding SHIP2, lead to Opsismodysplasia, a severe chondrodysplasia characterized by a shortening of the long bones and a delay in bone mineralization. Consistent with the human disease phenotype is the finding presented here that inhibition or deletion of SHIP2 reduces mineralization in chondrocyte and osteoblast models of mineralization without affecting cellular differentiation. When SHIP2 levels are reduced, more focal adhesions form and cell adhesion is increased. Furthermore, the results demonstrate that SHIP2 regulates mineralization via the actin cytoskeleton indirectly through the inhibition of RhoA/ROCK pathway. This data represents the first mechanistic link between SHIP2 and the RhoA/ROCK pathway in mineralizing cells.

Since inhibition of ROCK activity restores mineralization in SHIP2-negative cells, the data suggests that ROCK is overactive when SHIP2 activity is reduced leading to increased gene transcription and actin polymerization. Since the SHIP2 catalytic inhibitor displays the same effect on mineralization as the depletion of SHIP2, it is hypothesized that the effect of SHIP2 on RhoA/ ROCK is mediated through its catalytic activity. ROCK1/2 contains a pleckstrin homology (PH) domain that acts as a regulatory unit (37) and binds the ROCK catalytic domain thereby serving to inhibit ROCK activity. Thus, the binding of a ligand to the PH domain activates ROCK by liberating the catalytic domain (38,39). This is relevant for SHIP2 because while the ROCK PH domain can bind different phosphoinositides including PI(3,4)P2, only the SHIP2 substrate, PI(3,4,5)P3, has been shown to activate ROCK (40). An inhibitor of PI3-kinase inhibits ROCK activity *in vitro* confirming the importance of PI(3,4,5)P3 for ROCK activity (40).

It is further hypothesized that SHIP2 regulates ROCK activity in chondrocytes and osteoblasts through regulation of the level of PI(3,4,5)P3. The increased levels of PI(3,4,5)P3 due to the loss or inhibition of SHIP2 overstimulates ROCK activity leading to increased ECM production and reduced mineralization. A summary of this hypothesis is presented in Fig. 7.

**Figure 7:**
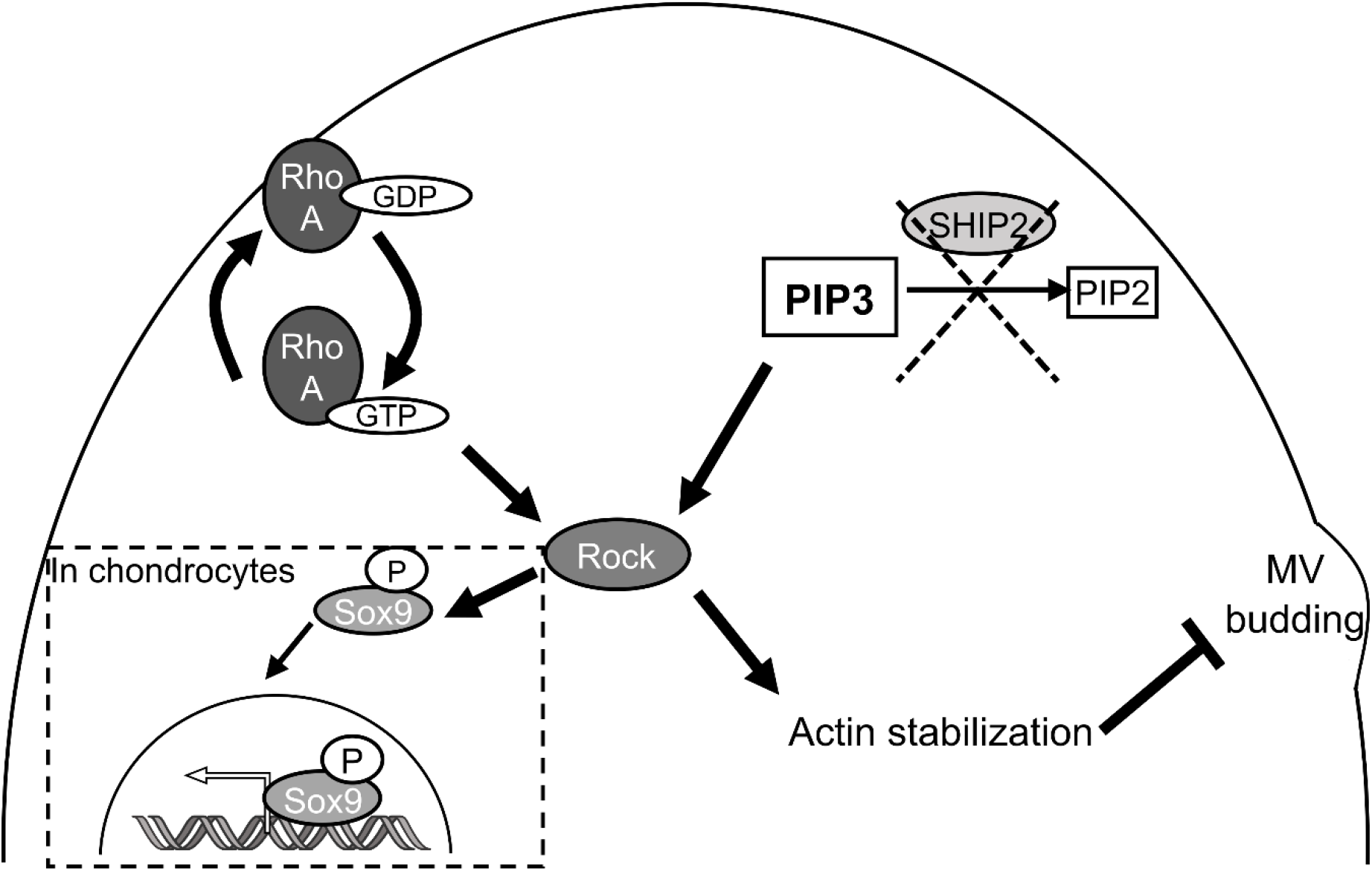
Hypothesis. In the absence of SHIP2, PI(3,4,5)P3 accumulates, activates ROCK which stimulates the production of ECM through the phosphorylation of Sox9 in chondrocytes. Increased ROCK activity also leads to stabilization of the actin cytoskeleton and inhibition of MV budding and ECM mineralization.

Through its main function, SHIP2 is considered as a negative regulator of the Akt pathway in insulin regulation. Akt has a PH domain and can bind both PI(3,4)P2 and PI(3,4,5). After recruitment, Akt activity is promoted by phosphorylation. Akt signaling can impact many different cellular processes. The data in the current study and those of others show that SHIP2 does not regulate Akt phosphorylation in chondrocytes (12). This result is consistent with reports showing that the absence of Akt in mice leads to osteopenia (43).

Several mouse models for SHIP2-inactivation have been generated, but the majority of these focus on the role of SHIP2 in metabolic processes, and only one studied the effect of loss of SHIP2 on skeletal development (9,11,12). The bone phenotype was relatively mild although it is clear that inhibition or absence of SHIP2 in chondrocytes reduced mineralization *in vivo* and *in vitro,* it also decreased chondrogenic marker and matrix marker expression without any effect on matrix synthesis. It is reported that SHIP2 regulates Erk signaling in different cell types. In rat chondrosarcoma cells, SHIP2 was shown to interact with several FGFR3 by mass spectrometry (44). In 293T and U2OS cells, SHIP2 interacted with FGFR1 and FGFR3 by immunoprecipitation leading to enhanced Erk signaling. This study shows a role for SHIP2 independently of its catalytic activity, instead SHIP2 acted as a scaffold that recruited Src family kinases to FGFR complexes (45). Despite significant efforts, an interaction between SHIP2 and FGFR1 or FGFR3 by immunoprecipitation in our model was not detected (data not shown). This discrepancy might be in part explained by the fact that Fafilek et al. used genetically-modified cells expressing tagged SHIP2 and tagged FGFRs to show the interaction thus increasing the level of protein expression. In the present studies, the level of protein expression might be too low to observe any interaction. Vande Catsyne et al., report that SHIP2 regulates mineralization in part through regulation of MEK and Erk1/2 upon stimulation by IGF1 but not by FGF2 in primary chondrocytes and ATDC5 cells (12). Erk1/2 levels were not altered in the ATDC5 model upon stimulation by IGF1 or LPA (not shown) but it is noted that primary cells and ATDC5 cells in a 3D micromass culture models were used in Vande Catsyne’s study, while the present study employed a 2D culture system.

The data show that in the absence of SHIP2, chondrocytes produce more proteoglycans. This was confirmed by an increase of the expression of several key ECM genes at the RNA level and is likely driven by increased Sox9 expression, a master regulator of ECM production in chondrocytes. It is reported that ROCK transcriptionally inhibits Sox9 but that inhibition does not necessarily affect Sox9 target gene expression, suggesting that posttranscriptional modifications might be important for Sox9 regulation (26,46). A 2010 study found that ROCK is able to directly phosphorylate Sox9 on Ser181, leading to the translocation to the nucleus necessary for its action (27). This is in agreement with the study findings, and further supports the hypothesis that ROCK is overactive in the SHIP2-negative ATDC5 model and would explain the overexpression of key Sox9 target genes observed in this study.

*Mmp13* is also overexpressed in SHIP2-negative ATDC5 cells; this was confirmed at the protein level. Increased MMP13 may be a response to the excessive production of matrix in these cells since MMP13 functions as a protease to cleave fibrillar collagens and aggrecan. RhoA/ROCK pathway has been shown to activate MMP13 expression in articular cartilage after compression stress (47), which could also correlate with the high expression of MMP13 that is seen in the ATDC5 model presented here.

The finding that an actin polymerization inhibitor rescues the mineralization phenotype in cells deleted for SHIP2 suggests a role for SHIP2 in formation of the actin cytoskeleton. This is supported by data showing increased cell adhesion and vinculin-positive loci present in cells deleted for SHIP2. It has been demonstrated in chondrocytes and in SaOs-2 cells that modulation of the actin cytoskeleton is involved in the release of matrix vesicles (MV) (48,49). MV are small vesicles released from the plasma membrane of mineralizing cells such as chondrocytes during endochondral ossification or osteoblasts during intramembranous ossification. MVs are engaged at the early steps of mineralization providing an optimal environment for the nucleation process of hydroxyapatite from Ca2+ and Pi (50). Phalloidin which stabilizes actin, inhibits MV release while cytochalasin D which depolymerizes actin enhances it. It is speculated that by indirectly inhibiting ROCK, SHIP2 contributes to the actin depolymerization that is necessary for MV release (Fig. 7).

In conclusion, this work highlights the role SHIP2 and the phosphatidylinositols play in osteoblast and chondrocyte ECM mineralization and, with the novel identification of the RhoA/ROCK pathway and actin remodeling in this process, potentially reveals new targets for the treatment of mineralization disorders.

## Supporting information

This article contains supporting information.

## Acknowledgements

Authors would like to thank Dr Gary Gibson, Dr Maozhou Yang and Jamie Endicott for helpful discussions and reviews.

